# BaYaka mothers balance childcare and subsistence tasks during collaborative foraging in Congo Basin

**DOI:** 10.1101/2024.01.22.576450

**Authors:** Amandine E. S. Visine, Adam H. Boyette, Yann Reische Ouamba, Sheina Lew-Levy, Mallika Sarma, Haneul Jang

## Abstract

Across cultures, mothers face trade-offs between childcare and other labor. In hunter-gatherer societies, mothers face this choice on a daily basis when deciding either to take infants on foraging trips or to leave them with caregivers in the village. Yet, it remains unclear how the presence of infants in foraging groups constrains mothers’ mobility during foraging. Here, we present GPS, energy expenditure and food returns data of 359 foraging trips of 23 BaYaka mothers in the Republic of the Congo. We find that mothers spent more time on out-of-village foraging activities when they took infants along, compared to when they left infants behind. However, infant presence in foraging groups does not affect mothers’ travel distance, travel range, energy expenditure or food returns. Regardless of infant presence, women travel longer and further in a larger area when foraging in groups, compared to when foraging alone, especially in groups with more adults, females and both kin and non-kin. Our results suggest that BaYaka mothers develop ways to accommodate childcare with foraging activities by combining individual-level and group-level behavioural strategies. Our study highlights that group foraging may allow mothers with infants to maintain high mobility, which may have been a key to human range expansion.

## Introduction

Humans have unique life-history traits including higher fertility, shorter inter-birth intervals, earlier weaning and an extended childhood period [1, 2, 3, 4]. Due to these traits, mothers face trade-offs in allocations of their investment per child [5, 6, 7, 8, 9], as well as of their time and energy between childcare and other labour [10, 11, 12, 13, 14, 15, 16]. In hunter-gatherer societies where individuals subsist via foraging, balancing the trade-offs between childcare and foraging could be a particularly critical problem for mothers [17, 18]. For example, mothers with young infants tend to reduce time allocated towards food acquisition to increase their time spent in infant care [13, 17]. This strategy can, however, result in increased risk to older offspring who still rely on mothers’ food provisioning. A wealth of literature in anthropology and human behaviour ecology shows that assistance from others can mitigate these trade-offs for mothers through direct childcare support and food provisioning [19, 20, 21, 22, 23, 24, 25, 26, 27, 28, 29]. Not all types of labour, however, are incompatible with childcare [30], as hunter-gatherer mothers often take infants along on foraging trips [13, 17, 18]. Yet, it is less clear what reproductive women with breastfeeding infants pay as the costs of childcare during foraging activities, particularly in terms of their mobility and travel ranges, and how the mothers balance these trade-offs. Considering the important role of women’s subsistence activities in family provisioning, exploring this topic in-depth can provide insights into constraints and opportunities for the remarkable demographic and range expansion characteristic of our species.

Although gathering activities are more compatible with childcare than hunting [31], studies showed that mothers with young children adjust their foraging activities by reducing foraging duration [13] or by engaging in low-risk, low-intensity activities [32, 33, 34]. Carrying infants is energetically expensive [35, 36, 37], especially for mothers who are still recovering from a long pregnancy and are using energy for breastfeeding [38]. When mothers take their infants along, therefore, they may have to use more calories to carry and breastfeed infants during the foraging trips, which requires higher energetic expenditure, compared to when they go foraging without infants. This energy drain for childcare may reduce maternal energy that could instead be put towards subsistence activities, as well as constrain mothers from travelling further away from the village and from exploring in a larger range. In addition, childcare—e.g., holding and breastfeeding in between foraging activities—may distract the mother from work, which further reduces the food acquisition rate [13, 18]. On the other hand, mothers have another choice of leaving infants with caregivers in the settlement (e.g., camp or village) [17], as humans are central place foragers—who return to a specific location after foraging trips to feed offspring and rest [39, 40]—and cooperative breeders—who receive childcare support from other group members [22, 41]. When mothers go foraging without their infants, mothers may save their energy from infant carrying and, thus, use fewer calories to walk, allowing them to have greater mobility, compared to foraging trips with their infants. In addition, mothers may have a surplus of energy available to spend on foraging activities, when they are free from the burden of infant carrying and breastfeeding. Nevertheless, mothers can have another constraining factor on their mobility as they have to come back earlier to the settlement to breastfeed their infants [13]. Therefore, mothers with breastfeeding infants face dilemmas in their daily decision-making of either taking or leaving infants.

Under the pooled energy model, such dilemmas can be mitigated by childcare support from allomaternal caregivers [17, 19]. Empirical studies show that allomaternal caregivers allow mothers to increase time allocation for other labour, reduce maternal energy expenditure, or maintain their baseline activity levels [17, 20, 14, 42]. Crucially, considering that women in different reproductive stages—e.g., mothers with breastfeeding infants versus mothers with weaned children—face different constraints [13], they have different need for childcare support. For example, as weaned children get much heavier to carry and are less risky to leave away from mother’s care, mothers with weaned children are more likely to leave children at the settlement, compared to mothers with breastfeeding infants [13], highlighting the important role of caregivers at the settlement. Compared to this, mothers with breastfeeding infants cannot leave their infants too long away from their care, because infants need to be breastfed. Hence, mothers with breastfeeding infants may take their infants along on foraging trips more often than mothers with weaned children, so that they can breastfeed infants intermittently in-between foraging activities. In this case, helpers in foraging groups could be much more important, and can provide childcare while mothers are acquiring food [18]. Supporting this, an empirical study showed that BaYaka mothers with infants have increased foraging efficiency when there are caregivers in foraging groups [18]. Nevertheless, other research shows that mothers with breastfeeding infants spend less time foraging and acquire less food in general than mothers with weaned children [13], indicating greater behavioural constraints in foraging behaviours of mothers with breastfeeding infants. Despite this possibly greater burden on mothers with breastfeeding infants compared to that of those mothers with weaned children, we lack detailed knowledge of how childcare responsibilities limit the nature or extent of mobility and foraging behaviour of mothers who are still breastfeeding their infants—beyond foraging duration and food returns—during their daily subsistence activities, and how these costs are mitigated by other individuals in foraging groups.

In this study, we aim to investigate in-depth how the presence versus absence of breastfeeding infants during foraging trips affects the mobility and foraging behaviours of mothers, by examining six different variables—travel duration, total travel distance, maximum distance from the village, foraging range, net energy expenditure and net food returns. For this, we present GPS tracks, heart rate, and food returns data of 359 foraging trips of 23 mothers who are breastfeeding their infants, from a BaYaka community comprised with 23 households with 180 individuals, in northern the Republic of the Congo. The BaYaka are a group of contemporary hunter-gatherers who practice hunting, gathering, fishing, and crop cultivation. Both BaYaka men and women engage in intense physical activity, but women spend more time in more intense physical activity than men [43]. BaYaka women habitually travel long distances for food acquisition in variable-sized and mixed-age groups including both adults and children, as well as in groups including both kin and non-kin [44]. BaYaka mothers face choices of either taking infants on foraging trips or leaving them with caregivers on a daily basis [18]. However, childcare is widely shared among the BaYaka, through kinship and reciprocity, alleviating mothers’ burden [14, 45, 18, 46]. Using daily observational data of 23 BaYaka mothers’ foraging group composition including the presence of mothers’ infants, we investigate how travel duration, total travel distance, maximum distance from the village, foraging range, net energy expenditure and net food returns of mothers compare 1) between foraging trips with versus without breastfeeding infants and 2) between group foraging versus solo foraging. We further examine the effects of age-classes, gender and genetic relatedness of group members. Due to the energetic burden of childcare during foraging trips, we expect that the presence of breastfeeding infants in foraging groups constrains mothers from travelling long distance, far away from the village and from exploring in larger range, requires mothers to spend higher energy expenditure and results in lower net food returns. However, BaYaka mothers do not generally travel alone, and they receive childcare support from other members in foraging groups (e.g., carrying or holding infants), which further buffers the cost of infant presence during foraging trips [18]. Hence, we expect that other individuals in foraging groups provide childcare support—either carrying or holding infants while mothers are walking and/or acquiring foods—and, thus, allows mothers to have greater mobility, save their energy, and higher net food returns, by receiving childcare support from group members. Understanding these dynamics can provide scope for the compatibility between childcare and subsistence tasks with the help of others, which may have been an important key to human demographic and ecological expansion.

## Results

Across the 359 foraging trips, 23 mothers took their infants on trips 58.7 percent of the time. BaYaka mothers went foraging mostly in a group with other individuals, and rarely went alone—only 14.7 % of 359 foraging trips. The foraging groups varied in size and composition (mean = 4.8, SD = 0.17, range: 1–21; see Supplementary Table 1 for details on each category). Across the 359 trips, BaYaka women went on mixed-activity foraging trips consisting of different activities: 290 trips of gathering (mushrooms, nuts, leaves or fruits), 177 trips of extracting wild yams, and 101 trips of hunting and fishing. In the subsections that follow, we outline the model results testing first, how the presence of infants in foraging groups affects mothers’ mobility and foraging outcomes—i.e., trip duration, total travel distance, maximum distance from the village, range area, energy expenditure, and food returns (See Supplementary Table 2 for descriptive statistics of these metrics), and second, how the interaction between the presence of infants and the presence of other individuals affects these outcome variables. To present our results, we evaluate the median of the posterior distributions for each parameter and its associated 95% credible intervals (*CI*).

### The effects of infant presence and other individuals on mothers’ foraging trips

First, we tested the effects of infant presence (yes/no) and the presence of other individuals in the group (yes/no) on the mothers’ foraging trip duration (hr), total travel distance (m), maximum distance from the village (m), foraging range (km2), energy expenditure (kcal/min), and food returns (kcal). We found that the presence of infants in foraging groups increased mothers’ travel duration, suggesting that BaYaka mothers have longer foraging trips when they took their infants on trips, compared to when they left their infants in the village (Figure 1a). However, the presence of infants in foraging groups did not have any considerable effects on mothers’ travel distance, travel range, energy expenditure or individual food returns (Figure 1b-f). Instead, we found that BaYaka mothers went on longer and farther foraging trips, and explored larger areas when they were accompanied by other group members, compared to when they went on foraging trips alone (Figure 1a-d). We found a weak tendency for mothers to have lower energy expenditure when they were foraging with other individuals, compared to when they were foraging alone (Figure 1e). However, the presence of other individuals during foraging activities did not have any considerable effects on mothers’ net food returns (Figure 1f)

**Figure 1.**
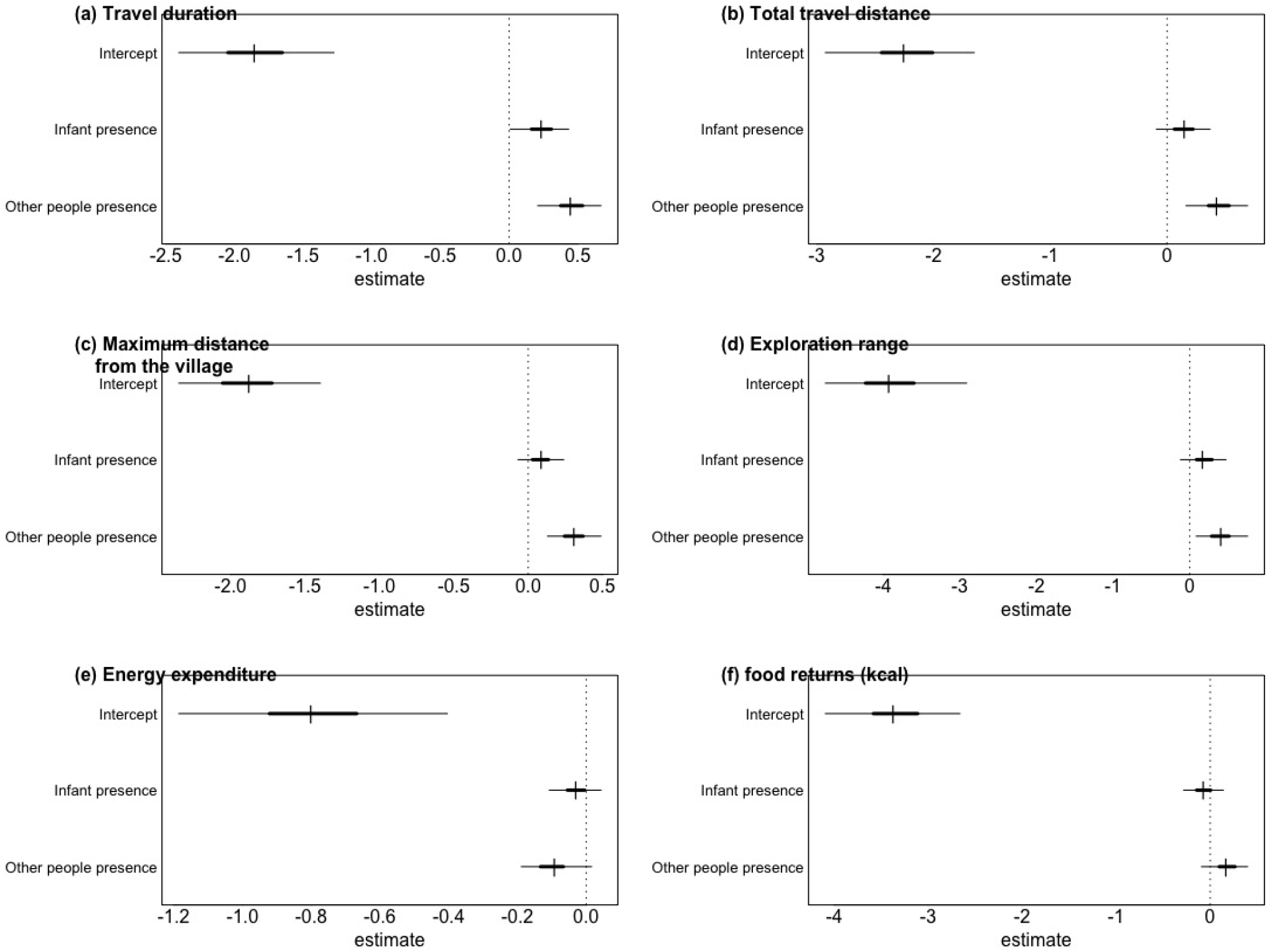
Forest plots showing estimated changes in (a) total travel duration (hr), (b) total travel distance (m), (c) maximum distance from the village (m), (d) range (km2), (e) energy expenditure (kcal/min) and (f) food returns (kcal) of focal mothers during foraging trips, according to the presence of the infant (yes/no) and the presence of any other members (yes/no) in foraging groups. All results present the median of the posterior distributions for each parameter, 50 percent (inner) and 90 percent (outer) credible intervals (*CI*).

### The effects of different group members on mothers’ foraging trips

We further aimed to understand which group members affected mothers’ travel duration, total distance, maximum distance, range size, energy expenditure, and food returns. For this, we investigated the interactive effects of infant presence and the number of group members in different categories—age-classes (adults/children), gender (females/males) and kinship relation with the focal mothers (kin/non-kin)—on these trip characteristics. We defined kin as when the genetic relatedness (r) of two individuals was between 0.125 and 0.5. Three sub-models were built to investigate the effects of age classes and gender of group members and their genetic relatedness with the focal mothers, respectively, on each outcome variable, resulting in a total of 18 models. We did not find any considerable effects of interactions between infant presence and the number of other group members of different categories (figure–forest plot in SI). Therefore, we dropped the interactions from each model and investigated the effects of each parameter, which we explain below in the order of different categories: age-classes (adults/children), gender (females/males) and kinship (kin/non-kin). First, we found that the number of adults (defined as 20 years and over) in groups increased mothers’ foraging trip duration, total travel distance, maximum distance from the village and exploration range (Figure 2a-d, first column). We did not find considerable effects of the number of either adults or children in foraging groups on maternal energy expenditure. The number of children (between 4 and 19 years), but not adults, tends to increase mothers’ individual food returns (Figure 2f, first column). Second, the number of females, but not males, in foraging groups increases the trip duration, total travel distance, maximum distance, and foraging range (Figure 2a-d, second column). Third, both kin and non-kin in foraging groups tend to increase BaYaka mothers’ trip duration, total travel distance, maximum distance, and foraging range (Figure 2a-d, third column). We found a weak tendency of kin effects on decreasing maternal energy expenditure, but not on food returns (Figure 2e-f).

**Figure 2.**
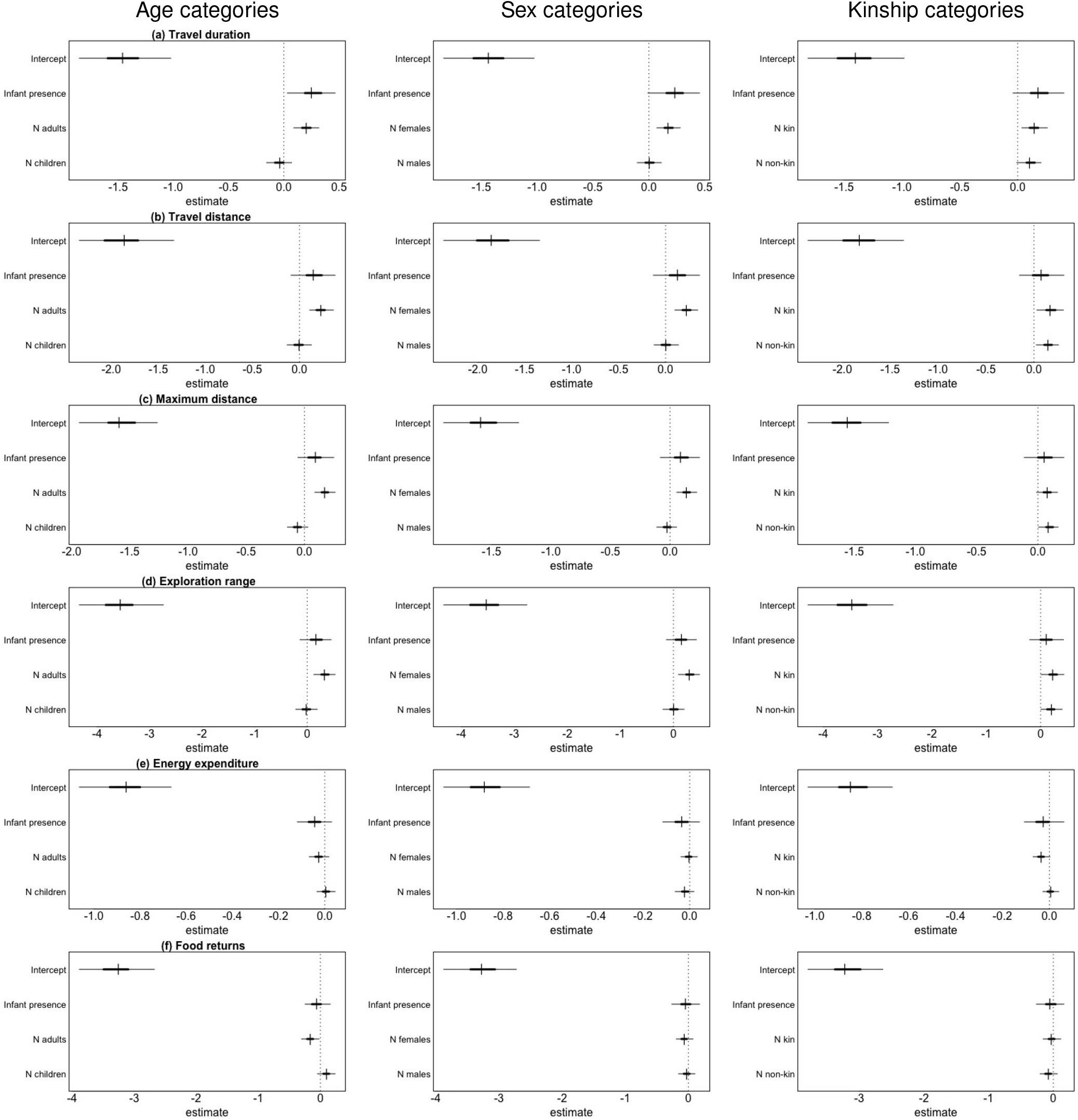
Forest plots showing estimated changes in (a) total travel duration (hr), (b) total travel distance (m), (c) maximum distance from the village (m), (d) range (km2), (e) energy expenditure (kcal/min) and (f) food returns (kcal) of focal mothers during foraging trips, according to the presence of the infant (yes/no) and the number of individuals in each category in foraging groups: age class (the first column), gender (the second column) and kinship (the third column). All results present the median of the posterior distributions for each parameter, 50 percent (inner) and 90 percent (outer) credible intervals (*CI*).

## Discussion

We have found that taking infants along on foraging trips does not decrease mothers’ mobility or net food returns. Regardless of infant presence, group foraging allows the mothers to have greater mobility compared to solo foraging. Our results suggest that BaYaka mothers accommodate childcare with foraging labour during collaborative foraging. Below, we discuss our results on the effects of infant presence during foraging trips and then the effects of group members.

We found that there were no considerable differences in mother’s total travel distance, maximum travel distance from the village or foraging range, nor in their net energy expenditure or net food returns, when mothers go foraging with versus without infants. These results can be explained by the confounding effects of individual-level constraints and opportunities, both when taking infants along and when leaving infants behind. When taking infants on foraging trips, the energetic burden of carrying and breastfeeding infants would restrict mothers’ mobility, but even without infants, mothers are constrained from travelling far away from the village, as they need to return to the village to breastfeed. Consistent with the findings from Hiwi foragers in western Venezuela [13], however, we found that BaYaka mothers spend more time on out-of-village foraging trips when they take their breastfeeding infants along, compared to when they leave them behind with caregivers in the settlement. When taking infants on foraging trips, the mothers can breastfeed infants intermittently during foraging activities without needing to return to the village and, thus, mothers can spend more time on foraging trips. Yet, longer foraging trip duration does not necessarily mean that mothers walk longer distances or spend more energy. Instead, we observed that the BaYaka mothers often spend time sitting or participating in less intense activities when they are with their breastfeeding infants, which was also shown in previous studies—mothers with young children tend to engage in low-risk, low-intensity activities [32, 33, 34]. In contrast, we observed that mothers engage more in high-intensity activities, such as cutting tree roots, digging ground, or carrying heavy baskets, when they go on foraging trips without infants. Therefore, our results and observations suggest that mothers who take infants on foraging trips spend longer time at a foraging patch, instead of exploring a large foraging range. This is also consistent with the finding from Hiwi mothers who stayed longer at a foraging patch when they were with their infants [13]. Although mothers’ foraging efficiency gets lower when they take their infants on foraging trips [18], mothers increase the total time spent on foraging activities to maintain the net food returns of the trips. Hence, this strategy allows mothers not only to compensate time spent for childcare during foraging, but also to collect sufficient food, without spending more energy walking while carrying infants as well as without the risk of harming infants by taking them to far-away, unfamiliar areas.

In addition to this individual-level strategy, we have found that BaYaka mothers go on longer and farther foraging trips and explore in a larger range when foraging in groups, compared to when foraging alone. Although the effects are weak, we also found that mothers have lower energy expenditure, but higher food returns, when they are accompanied by other group members. As in Figure **??**, we observed that when BaYaka mothers go foraging in a group, individuals in the foraging group help the mothers carry their infants and/or baskets, and thus, the mothers rarely carry both their infants and baskets. Hence, the help from group members allows mothers to travel longer distances in a large range, but also to maintain or reduce their overall energy expenditure. These results are consistent with the finding that Shuar women maintain their baseline activity levels through increased contributions from family members [42].

When we further took into account different categories (age class, sex, and kinship) of group members, we found that adults and females in foraging groups increase the mothers’ foraging trip duration and mobility. The presence of independent children, who can walk on their own and join women’s foraging trips, does not restrict women from travelling long distances in a larger range, which provides evidence that even young children can achieve adult-level walking capacity [47]. Crucially, consistent with the previous finding from Jang et al. [18], we found that children in foraging groups, but not other adults, are likely to increase the net food returns of the mothers, highlighting the role of children as helpers during women’s foraging trips. We observed that BaYaka children who accompany women’s foraging trips often hold and monitor infants when the mothers are collecting food sources, but also actively collect food items and put the collected food items in women’s baskets when the mothers stop foraging to take care of the infant or to rest. Compared to children, however, most adults put the collected food items in their own baskets during the resource harvest, resulting in no addition on the net food returns of the mothers, although women share foods once back in the village. In addition, we found that both kin and non-kin in foraging groups increase the mothers’ mobility, although kin tends to decrease maternal energy expenditure, suggesting that kin provide more physical help to the mothers during foraging trips (e.g., carrying heavy baskets or infants). The absence of kin bias and the positive effects of both kin and nonkin on women’s mobility suggest benefits to mixed-kinship foraging groups, possibley because they allow women to access a greater pool of resources and environmental knowledge [48] and, thus, to explore in larger ranges beyond what an individual or only-close kin group can. These findings highlight that women are able to maintain their net energy expenditure and net food returns with the help of other individuals that buffer maternal increased needs, and that women can extend their foraging range using collective knowledge. This provides a scope for the important role of group foraging in collective actions of foraging and childcare, as well as in mobility.

In summary, using the fine-grained data on daily foraging trips of BaYaka mothers, the present study investigated how taking breastfeeding infants along on foraging trips affects the mothers’ mobility, energetics, and foraging outcomes, and how foraging group members affect those relationships. Our study delivers important insights into the compatibility between childcare and subsistence work, specifically through mothers’ combining individual-level and group-level strategies. To better understand behavioural strategies, future studies need to take into account mothers’ actual time spent carrying infants versus foraging, for example, by conducting focal-follows. Moreover, comparing mobility between mothers with breastfeeding infants and mothers with weaned children will provide further insights into the extent to which reproductive-age women can combine their childcare and subsistence responsibilities, as empirical studies have so far only compared their foraging duration and food returns [13]. Furthermore, we observed that even when the mothers go on foraging trips without their own infants, they provide childcare for other women in foraging groups by carrying other infants, which might have diminished the effects of the presence of their own infants. Hence, to better examine the cost of infant presence at a group level, future studies should investigate how the presence of an infant as well as the total number of infants affect the total energy expenditure and food returns of all group members in a foraging group as one unit. This will allow us to understand the cost of childcare at a group level beyond the mothers’ individual level, as well as the group-level strategies to balance trade-offs between childcare and foraging, under the pooled energy model [19]. Nevertheless, our study provides empirical evidence for the important role of women’s group foraging by showing that foraging in groups allows mothers to travel in large ranges with little impact of carrying infants on their energetic costs or food returns. Our study helps to initiate future work that will elucidate the links between childcare and the physical well-being and/or constraints of mothers as well as mother’s strategies to mitigate these trade-offs, which is important to understand the demographic success of humans as well as human range expansion.

## Methods

### Ethics and Consent Procedures

Permissions to conduct this research in the Republic of the Congo were obtained from the Institut de Recherche en Sciences Exactes et Naturelles (IRSEN) and the Institut National de Recherche en Sciences Sociales et Humaines (INRSSH) in Brazzaville. H.J. presented an overview of the project including the research aims and methods in a public meeting with all residents in the village, and obtained the consent of the community. In the following days, H.J. visited each household and obtained individual and familial consent from focal mothers and her family members, through which all of them agreed to participate in daily interviews on their foraging trips and to wear GPS watches during foraging trips within the framework of this study.

### Data collection

The study was conducted between August and October 2022 in a BaYaka village in the forest of the Likouala department in the northern Republic of the Congo. In the beginning of the study period, H.J. conducted the household survey for each house in the village and collected demographic and household-level data for each family unit. All of 23 households in the study village were surveyed, and the village census counted 180 individuals. All 23 nursing mothers of the village, who have an infant, whose age ranges from 3 months to 3.5 years, and are involved in daily foraging trips were selected as focal women for our study. H.J. and Y.R.O. collected anthropometric data of all 180 individuals, namely height and weight, and estimated their ages and age classes based on the number of their children and siblings. During 6 weeks of the study period, we distributed 20 Forerunner 55 watches (Garmin, Olathe, United States) to focal mothers every morning and collected them in the evening on the same day. The GPS watches record GPS coordinates and heart rate measurements (in bpm) every second. We conducted daily short interviews with focal women when they left the village and when they returned from those trips by recording the time when they left the village and returned, trip types (e.g., foraging, crop cultivation, fetching water and firewood, social visits, etc.), the presence of their nursing infants in groups, and group member identities. We observed the group composition when they left and returned to the village, but we also asked the focal mothers all the names of the accompanying people to confirm it. When the focal mothers returned to the village with food sources, we recorded the types of the collected food items and measured the weight of each food item. In total, we recorded 359 person-days of GPS track with heart rate measurement and food return data from 23 focal mothers.

### Data processing

We first cleaned the GPS tracks by deleting the GPS points with unrealistic walking speeds that were higher than 30 km/h [49], which could have resulted from the errors of the satellite recording. Using the GPS coordinates of all the houses in the village, we defined the geographical centre of the village and the village area, and extracted GPS tracks only outside of the village area, to calculate metrics of out-of-village foraging trips. For each foraging trip, we calculated *total travel duration (hours), total travel distance (metres), maximum distance from the village (metres), range area (km2), net energy expenditure (kcal/minute)*, and *net caloric food returns (kcal)*. The ***total travel duration*** of the trips (hours) is the subtraction of the arrival and departure times. We calculated the ***total travel distance*** (metres) as the sum of the distances between each successive GPS waypoint of the foraging trip GPS tracks. The ***maximum distance from the village*** (metres) is calculated as the distance between the centre of the village and the furthest waypoint from the village centre. We calculated the ***range area*** (km2) by using the maximum convex polygon formula provided by the ‘mcp’ function in the ‘adehabitatHR’ package in R (version 0.4.20). For the ***net energy expenditure***, we used the method of Hiilloskorpi et al. [50], which has already been adapted for the BaYaka and allows us to calculate the energy expenditure using two different formula for intense activities (a heart rate above 90 bpm) and light activities (a heart rate below 90 bpm) [43]. We first calculated basal metabolic rate (BMR) for each focal woman, and then transformed heart rate (bpm) into energy expenditure (kcal/min), while taking into account the individual body weight (kg). Finally, we multiplied the mass (g) of each food item collected during a foraging trip by the nutritional value (kcal/g) from the Benoît et al. dataset [51], and then summed the caloric value (kcal) of all the collected food items to calculate the ***net caloric return*** of the foraging trip. All the data was processed in R (version 4.3.0) on Rstudio Pro (server version 2023.03.0).

### Statistical analyses

We used Bayesian multilevel regression models in the Stan computational framework (http://mc-stan.org/), accessed with the function ‘brm’ of the brms package v. 2.16.3 [52] in R v. 4.1.0 (R Core Team 2020). The unit of analysis in the statistical models was a subsistence trip of a focal woman, and we used six trip metrics of each foraging trip as response variables: *total travel duration (hours), total travel distance (metres), maximum distance from the village (metres), range area (km2), net energy expenditure (kcal/minute)*, and *net caloric food returns (kcal)*. To examine variation across focal women, days, and food items, we first fitted baseline models for each response outcome, which account for the multilevel structure of the data but include only random effects of focal women IDs (N = 23), day ID (N = 35), and targeted food types (N = 15 different combinations of 7 unique food items, including animals, mushrooms, leaves, nuts, tubers and fruits). Baseline models include no predictor variables (i.e., fixed effects). We checked the standard deviations of the log-odds for response outcomes between focal women, days and different food types, and calculated intra-class correlation coefficients to investigate the proportion of variance captured by each level (focal women ID, day ID, and food types) [53]. We then fitted models, including the fixed effects and random effects described above. To capture variation between focal women in infant presence, as well as foraging group composition, we included random slopes of the fixed effects within random intercepts of focal woman ID [54, 55].

First, we designed models with two fixed effects: the presence of the women’s infants (yes/no) and the presence of other individuals (yes/no) in foraging groups. To deal with between- and within-individual effects of focal women, we included both individual-level (focal woman-level) and trip-level fixed effects. As individual-level predictors, we calculated the average probability for each focal woman to take her infant on foraging trips and the average probability of going on foraging trips with other individuals. As trip-level predictors, we included the presence of a focal woman’s infant (yes/no) and the presence of other individuals (yes/no) during each trip. For the food returns model, we additionally included a predictor of a non-linear functional form between travel duration and food returns, as we assumed that food collection duration can affect food returns.

Second, we ran models to investigate the effects of group members’ age-classes (adults versus children), genders (females versus males), and kin relations to the focal women (kin versus non-kin), respectively. Specifically, we included two-way interactive effects between the presence of infants in groups (yes/no) and the number of individuals in each category. In the age-class effect model, we included two two-way interactions: 1) between the presence of infants and the number of adults, and 2) between the presence of infants and the number of older children in foraging groups. In the gender-effect model, we included interactions between infant presence and 1) the number of females and 2) the number of males. In the kinship-effect model, we included interactions between infant presence and 1) the number of kin (0.125 *≤* r *≤* 0.5) and 2) the number of non-kin. We used nonlinear functions for these interactions, using a spline. As individual-level predictors, we calculated the average probability of each focal woman to take her infant on foraging trips, as well as calculated the average number of other individuals who accompanied foraging groups of each focal woman. As trip-level predictors, we included infant presence (yes/no) and the number of individuals in each category for each trip with an interaction with infant presence. After we did not find any considerable effects of those two-way interactions, we ran the models with individual parameters after dropping interactions, to investigate the independent effects of each predictor.

All quantitative predictors were standardised to a mean of zero and a standard deviation of one before fitting models [56]. All models were fitted with a log-normal error distribution. We used weakly informative normal priors to guard against overfitting [57]. We obtained posterior distributions of the effects of predictors from four independent MCMC chains. We compared each of the main models to the baseline model with information criteria, using model_weights in the brms package. We found that the models with fixed effects were favored by model comparison.

## Acknowledgements

We thank the Ministère de la Recherche Scientifique et de l’Innovation Technologique, the Comité d’Ethique de la Recherche en Sciences de la Santé, Institut de Recherche en Sciences Exactes et Naturelles (IRSEN) and Institut National de Recherche en Sciences Sociales et Humaines (INRSSH) for their permission to conduct our research. We gratefully acknowledge Prof. G. Tchimbakala at IRSEN, Prof. G. Moussavou at INRSSH and V. Kandza for logistic support for this study. We thank E. Ringen for helpful advice for the statistical analyses. We are especially grateful to the BaYaka family for allowing us to follow them on their daily subsistence expeditions.

## Authors contributions

H.J. conceived the research; H.J. designed the data collection protocol with the help of A.H.B., S.L-L. and M.S.; H.J. and Y.R.O. collected the data; A.V. cleaned and prepared the data for statistical analyses under the supervision of H.J.; A.V. and H.J. conducted statistical analyses and interpreted the results; A.V. and H.J. wrote the paper; all authors reviewed and edited the paper.

## Data and material availability

All relevant data and code for reproducing the analyses and figures are available at the follow-ing GitHub repository: https://github.com/haneuljangkr/bayaka-tradeoffs-childcare-foraging

