## Supplementary for "BaYaka mothers balance childcare and subsistence tasks during collaborative foraging in Congo Basin"

### Data processing

All the data were processed and analysed in R (version 4.3.0) on Rstudio Pro (server version 2023.03.0). The focal women made an average of 23 subsistence trips each (rank: 8 - 44), and a total average of 14 subsistence trips per day (rank: 1 - 26). In total, we recorded 650 person-days of GPS tracks with heart rate measure, conducted 985 interviews when they left the village and 962 interviews when they returned to the village, and measured 598 food returns from 23 focal mothers. Those trips could be classified into different categories (*work for the Bantus*, *firewood*, *water* and *subsistence*), with different places visited during out of the village trips, e.g., firewood, water pond, neighbouring village, forest and/or garden. In order to include only the foraging trips in our analyses, we removed the trips to fetch water and firewood, and to visit other villages for social purposes, and only used the trips with the category of *subsistence*, which resulted in three places visited: *forest*, *garden* and *forest and garden*. In total, we had 359 subsistence trips for statistical analyses. 352 among 359 foraging trips have data on food returns (i.e., the sum of each food item's nutritional value in kcal, before it has been shared or processed).

All the people in the community were turned anonymous and, using the demographic and household data, a kinship grid was set up. Kinship was considered when proximity (r) was between 0.125 and 0.5. We recorded the sex (woman or men) and age (three classes: infant [0-3 years], children [4-19 years], and adult [20 years and over]) of each members of the community. We recorded the accompanying individuals of each subsistence trip of focal women, to determine the composition of the group. When unknown individuals (e.g., visitors) joined the subsistence trips, we asked and recorded their age, sex and their relatedness to the focal women.

GPS coordinates in degrees-minutes (example format: N2° 19.554', E16° 32.628'), have been converted into decimal degrees (example: Y = 2.326, X = 16.544). Using the GPS coordinates of all houses in the study village, we defined the geographical centre of the village, and the area of the village by drawing a circle whose radius is the distance between the geographical centre and the furthest house from the village centre. We added 5 metres to this distance and used it as the radius of the village area, in order to account for GPS movements around the house, rather than a subsistence trip outside the village. All GPS data outside the village was therefore kept. We removed gps tracks of the subsistence trips in which the last GPS waypoint of the trip . did not end in the village, but ended more than 30 metres from the edge of the village, because this means that the GPS tracks stopped recording and the data were not complete.

| Group members | Sex | Mean | Range | Std error |
|---|---|---|---|---|
| Infants | Both | 1.1 | 0 - 5 | 0.06 |
| (0 months to 3 years) | Female | 0.5 | 0 - 4 | 0.04 |
| | Male | 0.6 | 0 - 3 | 0.03 |
| Children | Both | 1.0 | 0 - 7 | 0.08 |
| (4 to 19 years) | Female | 0.6 | 0 - 4 | 0.04 |
| | Male | 0.6 | 0 - 3 | 0.03 |
| Adults | Both | 2.3 | 0 - 9 | 0.09 |
| (20 years and over) | Female | 2.2 | 0 - 9 | 0.09 |
| | Male | 0.1 | 0 - 3 | 0.02 |
| Kin | Both | 1.6 | 0 - 10 | 0.09 |
| ($0.5 \geq r \geq 0.125$) | Female | 1.0 | 0 - 9 | 0.07 |
| | Male | 0.7 | 0 - 6 | 0.04 |
| Non-kin | Both | 1.9 | 0 - 18 | 0.17 |
| | Female | 1.6 | 0 - 14 | 0.14 |
| | Male | 0.5 | 0 - 5 | 0.05 |

**Table 1.** Descriptive statistics of the foraging trip group composition for each age category (infants, children, adults) and genetic relationship (kin or non-kin) to the focal women.

| Trip descriptor | Mean | Range | Std error | Trips |
|---|---|---|---|---|
| Duration (hr) | 3.80 | 0.02 - 10.36 | 0.13 | 359 |
| Total distance (m) | 7355.90 | 10.04 - 26321.11 | 314.21 | 359 |
| Maximum distance (m) | 1425.60 | 200.00 - 5269.30 | 51.13 | 359 |
| Total area (km2) | 0.47 | 0.00 - 4.80 | 0.04 | 359 |
| Energy expenditure (kcal/min) | 2.07 | 0.84 - 4.52 | 0.03 | 359 |
| Caloric return (kcal) | 5070.94 | 40.02 - 73656.00 | 337.20 | 352 |

**Table 2.** Summary of the subsistence trip descriptors calculated for the statistical models.
